# A network module for the Perseus software for computational proteomics facilitates proteome interaction graph analysis

**DOI:** 10.1101/447268

**Authors:** Jan Rudolph, Cox Jürgen

**Affiliations:** Computational Systems Biochemistry, Max-Planck Institute of Biochemistry, Am Klopferspitz 18, 82152 Martinsried, Germany.

## Abstract

Proteomics data analysis strongly benefits from not studying single proteins in isolation but taking their multivariate interdependence into account. We introduce PerseusNet, the new Perseus network module for the biological analysis of proteomics data. Proteomics is commonly used to generate networks, e.g. with affinity purification experiments, but networks are also used to explore proteomics data. PerseusNet supports the biomedical researcher for both modes of data analysis with a multitude of activities. For affinity purification, a volcano plot-based statistical analysis method for network generation is featured which is scalable to large numbers of baits. For posttranslational modifications of proteins, such as phosphorylation, a collection of dedicated network analysis tools helps elucidating cellular signaling events. Co-expression network analysis of proteomics data adopts established tools from transcriptome co-expression analysis. PerseusNet is extensible through a plug-in architecture in a multi-lingual way, integrating analyses in C#, Python and R and is freely available at http://www.perseus-framework.org.

## INTRODUCTION

The study of complex systems^1^ is concerned with the question of how the relationship between the parts of a system give rise to its collective behavior. Complex systems often generate emergent properties^2^ which are not contained in an obvious way in its parts. Examples of such networks range over all disciplines of science, including the study of social media networks^3^, scientific collaboration networks^4^ and the human brain and its interconnected neurons as a particularly interesting one. The interactions between the components of a complex system define a network of connections consisting of nodes and edges. Much of the relevant content is concealed in the network constructed from these interactions and is not visible in the components themselves. For instance, the brain connectome^5^ is believed to make us who we are and not the cellular content of the brain^6^. Similarly, the observation of cellular concentrations of biomolecules without considering their interaction would provide a limited picture that ignores potential emergent properties of the biomolecular complex system. Hence it is mandatory to study biological systems, such as cellular concentrations of biomolecules, in the framework of network biology^7^.

At a fundamental level, all network connections between the cellular biomolecules are biochemical reactions and their specification in biochemical pathways together with their subcellular spatial distribution would provide complete knowledge about the biological network state of the cell. This collective network of all biochemical reactions contains all metabolic reactions, the signaling cascades, gene regulatory networks and all complex-forming non-covalent interactions between molecules, as for instance protein-protein interactions. Due to the limitations of experimental and computational methods to map out this interaction network, we often obtain only partial knowledge about the complete biochemical reaction network from experiments. Networks are however not limited to describing fundamental physicochemical interactions between biomolecules. For instance, in a gene co-expression network analysis^8^ one looks for similarity of expression patterns of gene products over many samples. Strongly correlated expression implies that these genes have some kind of non-physical interaction, e.g. they are part of the same transcriptional regulatory program or they share membership in the same pathway or protein complex. However, the exact relationship in terms of biochemical reactions remains unknown with these and other techniques. Hence, in these cases, networks describe a coarser grained level of detail, in which relationships between molecules are not necessarily biochemical reactions, but of a more general kind.

Computational proteomics is a mature data science that copes well with the large amounts of data produced in mass spectrometry experiments^9^. Perseus is an established framework for the downstream bioinformatics analysis of quantitative proteomics data^10,11^. The initial version of Perseus provided a comprehensive framework and set of activities to analyze data matrices originating from quantitative proteomics in a workflow environment. The main idea behind Perseus is to enable the researchers in biomedical sciences to perform the data analysis themselves. Here we describe how we extend this program to the analysis of biological networks in the context of proteomics. While cytoscape^12^ exists as the de-facto standard for network analysis and visualization, many proteomics-specific tasks for the generation and analysis of networks are lacking from this framework, as well as workflow navigation. PerseusNet fills this gap and enables non-computational experts to perform complete network-based analysis of their data. We explicitly do not want to re-invent existing methods and algorithms. Instead we designed an extensible framework that integrates with existing tools, like cytoscape, and interoperates with existing code and scripts from the network analysis community that were written in diverse languages, like Python and R. The data structures within Perseus that hold the networks were set up in a way that facilitates studying dynamic changes in networks and finding differential network properties over complex experimental designs. Side-by-side analysis of networks with data matrices in a common workflow environment allows for a seamless transition between matrix-centric and network-centric approaches.

In the following we start with a general description of the new network framework in Perseus, including how it enables multilingual programming and usage of code resources from R and Python. Then we introduce the new volcano-plot based analysis workflow scalable to large affinity purification-mass spectrometry (AP-MS) datasets. We describe how general and more specifically, large-scale protein-protein interaction (PPI) networks are handled and curated in Perseus. A section on the analysis of posttranslational modification (PTM) induced networks, like kinase-substrate relationships for phospho-proteomics is next. Finally, we cover co-expression analysis in Perseus and its applications to clinical proteomics.

## RESULTS

### WORKFLOW-BASED BIOLOGICAL NETWORK ANALYSIS

PerseusNet was devised to fulfill the computational needs of proteomics researchers wishing to accomplish network analysis of their data. While it is extensible through a new plugin application programming interface (API), and hence any network analysis functionality can be implemented, most tools needed for proteomics research and connecting it to generic network analysis platforms are included in the software (Fig. 1). Dedicated activities for analyzing AP-MS datasets and phospho-proteomics experiments in the context of kinase-substrate networks belong to the basic infrastructure of PerseusNet. The most common standard data formats are supported as input. An extended multi-language plugin API allows leveraging many existing tools in the analysis workflow. As an important example, co-expression clustering tools are integrated in this way.

**Figure 1.**
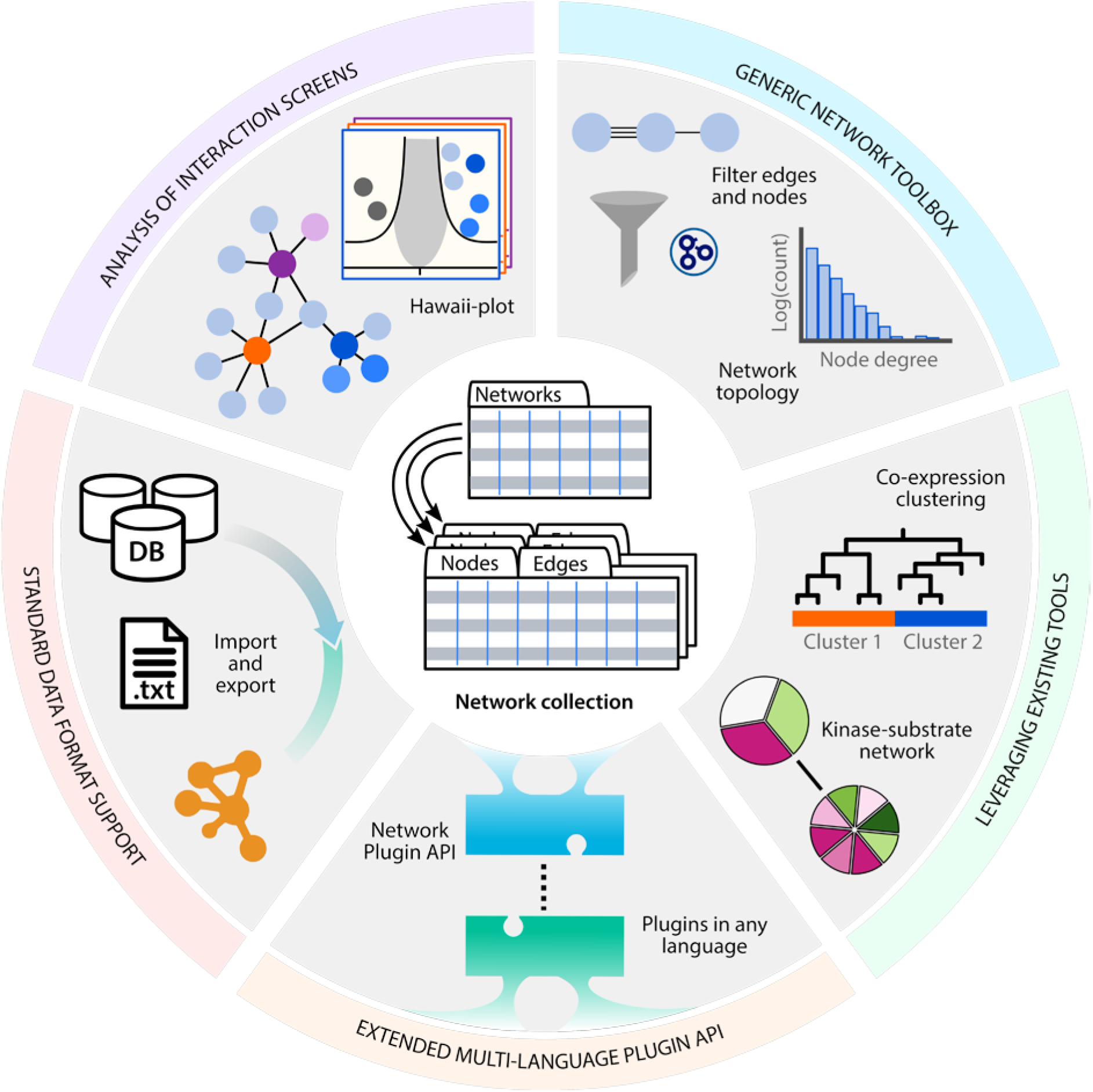
Schematic overview of the new network functionality in Perseus. PerseusNet implements a number of processing and analysis steps facilitated by the network collection data type. While including proteomics centric analyses, such as for the analysis of interaction screens, the network module also provides a number of general purpose tools, as, for instance, for network annotation, filtering, and topology determination. With the extension of the Perseus plugin API to networks and furthermore to other programing languages, it becomes possible to integrate existing network analysis tools in Perseus. Networks are easily imported to, and exported from Perseus, due to its support for standard formats.

To accommodate PerseusNet, we extended the Perseus framework with a new data type termed network collection (Fig. 2) that represents a set of one or more networks which are analyzed jointly in the workflow. Different networks within the same network collection can, for instance, represent networks derived from different individuals (patients), experimental conditions or biological replicates. All information in the network collection is organized in data tables, leveraging the existing augmented data matrix^10^ in Perseus. General information on the networks in the collection are stored in the networks table, where each row represents an individual network. Here, sample-related annotations, such as calculated global network properties, can be stored to enable their usage in analysis activities operating on a network collection. For instance, if the samples correspond to different patients, the networks table can hold patient-specific information as derived from patient records or questionnaires. These variables can then be used as independent or confounding factors in statistical analysis of the networks.

**Figure 2.**
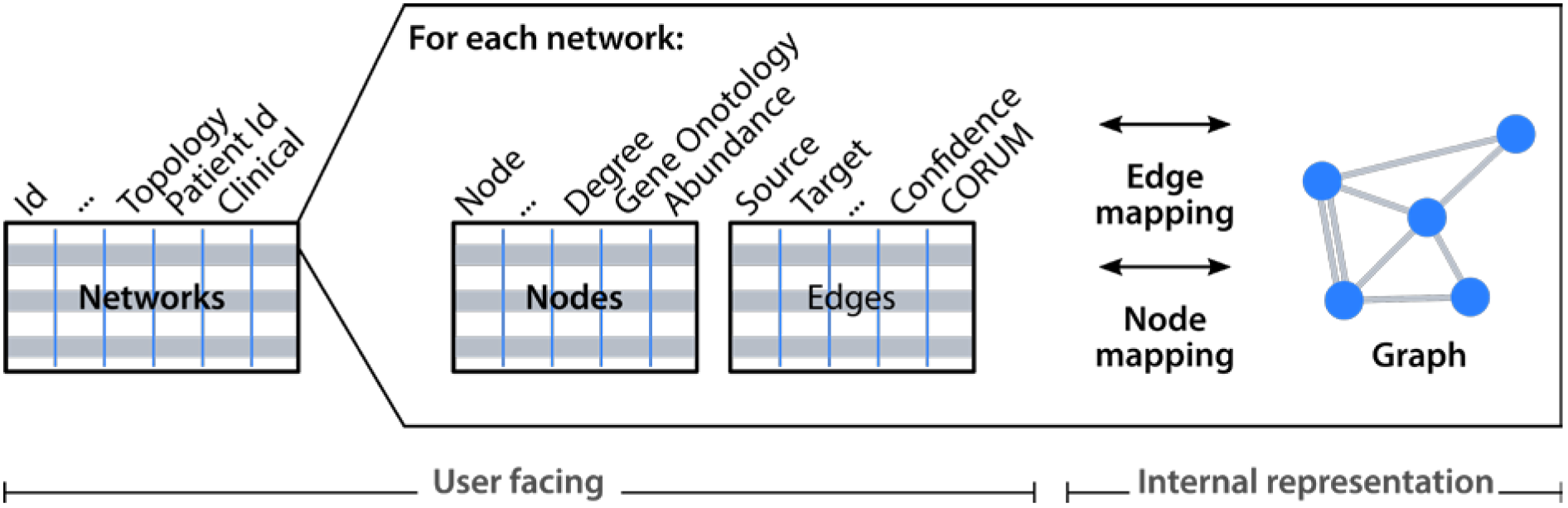
Schematic representation of the network collection data type. User facing information is displayed in tabular form with tables listing the networks in the collection, as well as providing detailed information on the nodes and edges of each network. Internally an auxiliary graph data structure aids in the implementation of graph algorithms. Node- and edge-mapping provide the required cross-references between the tabular and graph representation.

The nodes and edges of each individual network are stored in a pair of separate tables. The nodes table further describes the entities in the network, while the edges table provides details on the connections between the entities. The entities in the nodes table can be annotated with local network properties, such as the node degree. In case the entities correspond to proteins, biologically meaningful annotations could include membership in gene ontology terms, pathways or protein complexes. Similarly, edges can be annotated in the edges table with properties of pairwise relationships between proteins, as for instance interaction confidence measures. All of these properties are then accessible to the network analysis tools. Furthermore, all mentioned tables can be sorted and searched, allowing all information to be browsed and inspected intuitively. Internally, a graph data structure for each network enables the efficient execution of graph algorithms. We did not aim to include generic graphical representation of networks as node-link diagrams, since this can be achieved in other tools such as Cytoscape, for which we provide simple adaptors for the transfer of networks. However, several activities include specialized visualizations tailored to specific analyses.

In Perseus, all data analysis steps are performed within a graphical workflow. (See Supplementary Fig. 1.) Enabled by the newly implemented network collection, the Perseus workflow is now capable of all import, processing, and analysis steps in the side-by-side analysis of expression matrices and networks. All data that is imported into Perseus is represented as a separate entity in the workflow. Any matrix or network undergoing a processing step is not modified in-place but rather becomes a new entity that gets connected to the original data in the workflow. By inspecting both, input and output data, every step in the analysis is traceable and easily understood. Certain processing steps allow for the transformation of matrices into networks and vice-versa, or the mapping of data between the two. As a result, any analysis performed in Perseus, potentially including several side-by-side processing steps of networks and matrices, always remains transparent to the user.

### MULTILINGUAL PLUG-IN ACTIVITIES

The network collection data structure (Fig. 2) and the extended Perseus workflow provide the foundation for enabling various network analyses, many of which are available in Perseus. In general, networks either originate from external sources, or are created in a data-driven manner from within the workflow. To facilitate the import of external networks into the workflow, we implemented parsers for standard network formats, such as edge table (.tab|.txt|.csv), GraphML^1^ (.gml), Cytoscape’s simple interaction format^2^ (.sif) and D3js’s JSONgraph^3^ (.json) which enable loading interactions from most popular network databases, including STRING^13^, BioGRID^14^, IntAct^15^, CORUM^16^ and PhosphoSitePlus^17^. Furthermore, specific quantitative expression data, such as AP-MS drives the creation of novel protein-protein interaction networks, and phospho-proteomics datasets allow for a more detailed view or construction of kinase-substrate relationship networks. Specialized visualizations of such networks are provided (see later sections), which allow for an intuitive visual inspection of the results of the analysis. Perseus is not limited to physical interaction networks: co-expression clustering provides a powerful alternative to regular hierarchical clustering for expression proteomics studies. Finally, any network collection can be exported from the workflow in a plain text file format for sharing or use in any other external tools, such as Cytoscape. In order to accommodate these new capabilities in the Perseus plugin system, we extended the Perseus plugin API with new programming interfaces for the network collection and other associated data types, as well as the respective import, processing, and analysis interfaces. (See Supplementary Fig. 2.) This fully-featured API is available to all developers wishing to extend Perseus’ functionality with plugins. All analyses presented in this manuscript adhere to the new API.

In-order to better leverage the existing network analysis ecosystem, we additionally implemented a new mode of interoperability between Perseus and external tools (Fig. 3). The PluginInterop project enables this functionality, and allows the user to run external tools from within the Perseus workflow, most prominently scripts written in the popular R and Python languages. Open-source companion libraries for R^4^ and Python^5^ provide utilities for interfacing with Perseus. As a result, network analysis tools originally implemented in external tools can run from within the Perseus workflow with only minor adjustments. The implementations of the PHOTON and WGCNA plugins presented in the manuscript are based upon PluginInterop and its companion libraries. Instructions for interested developers on how to write scripts for Perseus or how to adapt existing tools can be found on the PluginInterop website^6^. In the following sections, we will present a number of network analyses which are now implemented in Perseus, with focus on their application to different types of proteomics data.

**Figure 3.**
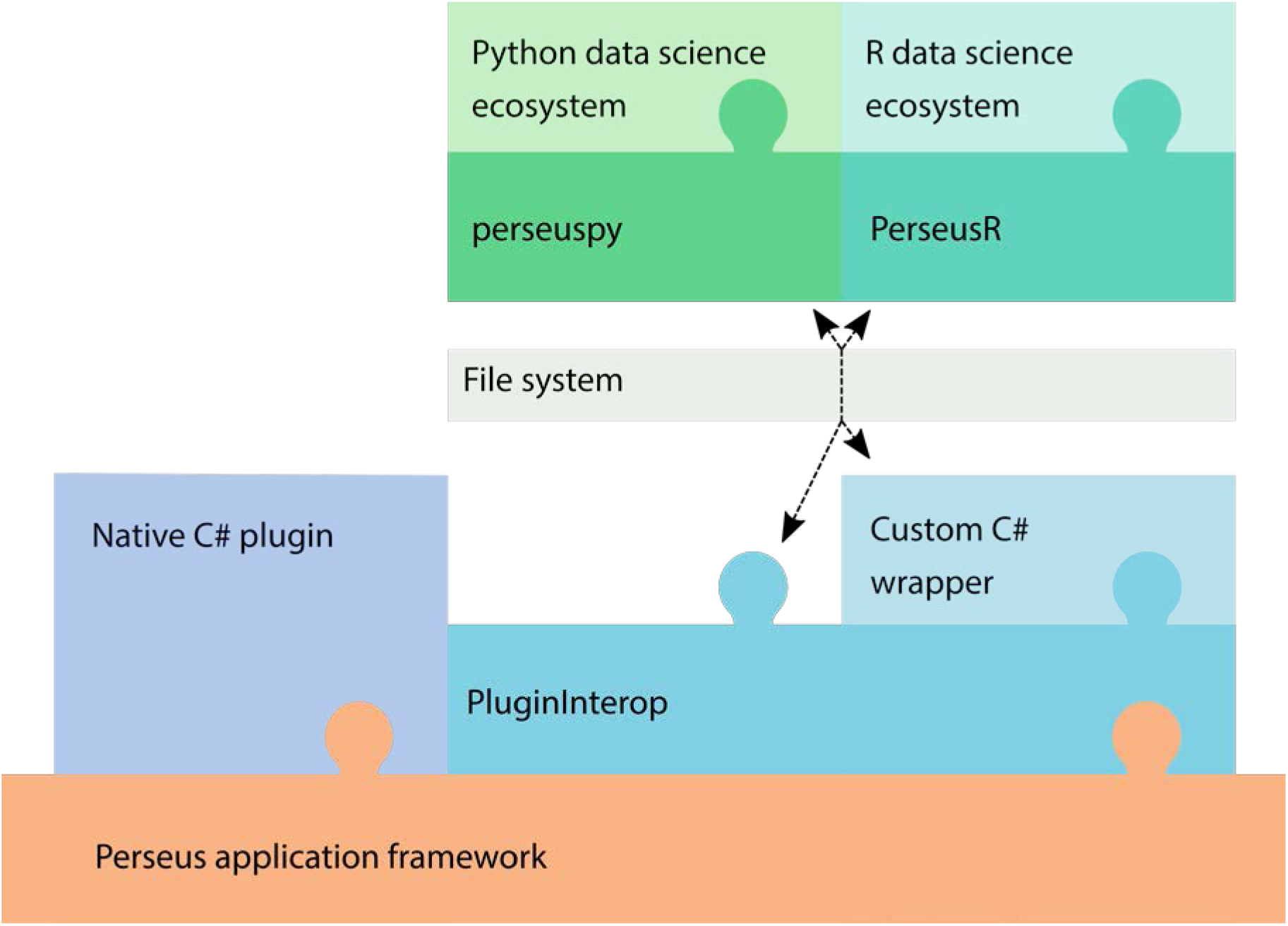
Schematic of the Perseus plugin system. Plugins written in C# are native to Perseus and implement their functionality directly on top of the APIs and data structures provided by the application framework. PluginInterop enables the execution of scripts in the Python and R languages, as well as other external programs. By communicating via the file system, data is transferred between Perseus and the external program. The companion libraries perseuspy and PerseusR enable developers to access the data science ecosystem in their language of choice. For custom GUI elements and an improved user experience of external tools, developers can implement a thin C# wrapper class that extends the generic functionality of PluginInterop.

### AFFINITY ENRICHMENT MS INTERACTOMICS

Affinity purification or enrichment coupled to MS analysis has become a powerful tool for interrogating PPIs^18,19^. It is able to provide not only a detailed view on proteins of interest, but it can also determine the basic building blocks for the assembly of large-scale protein-protein interaction networks^20,21^. Historically, protein complex members were detected by subjecting the sample to a series of purification steps followed by MS identification. With the advent of quantitative MS, detecting even transient interactions has become possible by relying not only on the identification itself, but instead on quantitative information. The sample is not purified, but only enriched for the protein of interest and its interaction partners and then subjected to MS quantification^22^.

Confidently identifying bona-fide interactions and distinguishing them from background binders, arising from off-target binding or contamination, requires data analysis of replicate case and control measurements. Compared to purely fold change-based methods, statistical tests provide a powerful way to compare case and control samples by calculating a test statistic and an associated p-value and limit the number of false-positives. For visual inspection of the results, the (negative logarithm of the) p-value can be plotted against the size of the effect, i.e. the difference between the means of logarithmic abundances, in a so-called volcano plot. Since one statistical test is performed for each protein, which amounts to a large number of tests performed simultaneously, the significance level needs to be adjusted to avoid increased numbers of false positives due to the multiple hypothesis testing problem^23^. A popular strategy to adjust for multiple testing is to control the false discovery rate (FDR), which can be achieved by permutation-based methods. Furthermore, in the volcano plot method it is necessary to define the functional form of the curves that separate significant from non-significant hits, either by straight lines, or in a more sophisticated way, introduced in the significance analysis of microarrays (SAM) method^24^, by modifying the t-test statistic with the background variance parameter s_0_. This standard workflow is available in Perseus but becomes increasingly cumbersome for interaction screens with more than a handful of baits. Parameter values for s_0_ and the FDR thresholds are often applied separately for each pulldown, inviting overfitting and cherry-picking, and also requiring results be subsequently combined manually.

We implemented the interactive multi-volcano plot (Fig. 4a) to analyze interaction screens with arbitrarily many baits and conditions simultaneously. Given the experimental design of the dataset, defined by baits and conditions, the analysis is applied to each experiment. For sufficiently large datasets, instead of dedicated control samples, an internal control can be assembled from the dataset for each pulldown consisting of pulldowns of other, unrelated baits. The results can be inspected through an interactive user interface. All volcano plots are displayed in the overview panel. A multi-functional detail panel shows more information on selected plots and provides zoom, protein selection and labeling options. If a single plot is selected, the volcano plot is shown in the detail panel. When two plots are selected, the t-test differences between the selected experiments are plotted against each other, highlighting changes in the enrichment of proteins between experiments (Fig. 4b). Additionally, all data can be browsed in tabular form, making it easily searchable and allowing for rich styling options. Known interactors or gene ontology annotations matching the experiment can be used to highlight proteins in the plot and can serve as a positive control for the adjustment of test parameters. All test parameters are controlled on a global level, effectively preventing overfitting and cherry-picking parameter values. We integrated the multi-volcano analysis into the new network module. Results from protein-protein interaction screens can be exported as network objects into the Perseus workflow. A specialized node-link visualization based on the open-source cytoscape.js library^25,26^ with multiple layers of information, allows for easy interpretation of the results (Fig. 4c). A protein-protein interaction network that was newly created in this way can be integrated with existing networks, or exported in various formats using the functions available through the network module.

**Figure 4.**
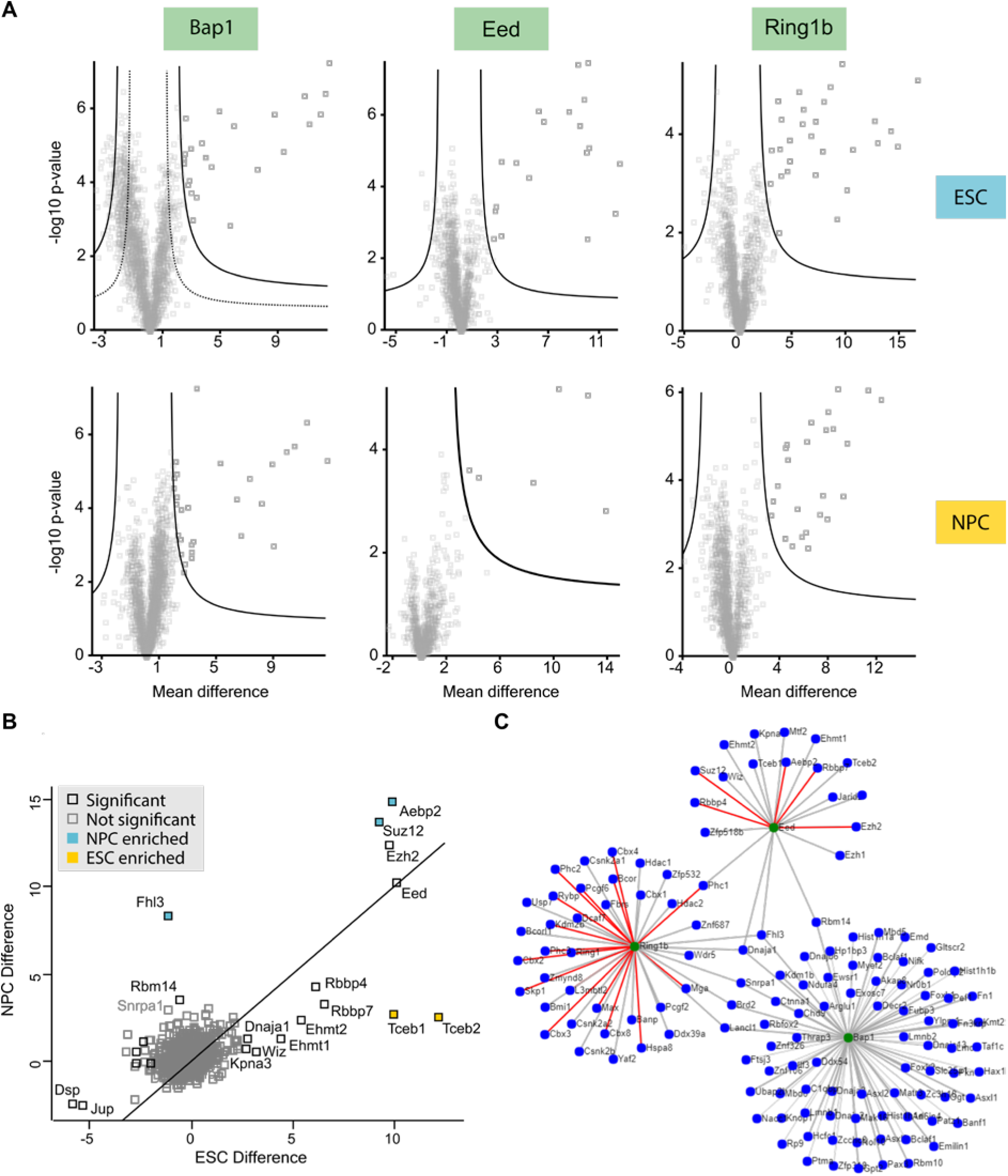
AP-MS. **a** The Hawaii plot provides an overview over the entire dataset27 (Supplementary Table 1 and online methods) consisting of three baits in two conditions. Significant interactors are determined using a permutation-based FDR and the resulting Class A (solid line) and Class B (dashed lines) thresholds are displayed in the plot. Class A interactors are displayed in dark grey, other proteins are shown in light grey. **b** Enrichment plot comparing the Eed pull downs in ESC and NPC cell lines. Significant interactors in any of the two conditions are displayed in black, non-significant proteins are displayed in light grey. Proteins differentially enriched in one of the two conditions will be located far from the diagonal and can be identified visually. **c** Visualization of the resulting protein interaction network for both cell lines. Bait proteins are colored in green and their interactors are colored in blue. Thick lines represent Class A interactions, thinner lines Class B. Interactions which were already annotated in the human CORUM database are highlighted in red.

As an example application, we obtained pull-down experiments from reference^27^, covering three baits in two different cell types. The filtered data set contained 2995 proteins. Using the new multi-volcano analysis (Fig. 4a), we obtained a PPI network with 134 nodes and 140 edges. The results were comparable to the original publication with overlaps between 55% (Ring1b ESC) and 91% (Bap1 ESC) for Class A interactions. Differences can be explained by the slightly different methodology used in this manuscript. We used the s0-modified t-test with s0 set to 1.0, and FDRs of 0.01% and 0.2% for Class A and B, respectively, while the authors of reference^27^ used individually chosen fold-change and p-value cutoffs for each experiment. Using the built-in visualization features, such as the enrichment between experiments, we identified several interactions that were conditional on the cell type (Fig. 4b). By annotating the newly created protein-interaction network with known complex interaction from CORUM and inspecting the resulting node-link network visualization (Fig. 4c), previously known and possibly novel interactions could be distinguished.

Further confidence in the existence of an interaction between a protein identified in a pulldown and the bait can be obtained by correlation analysis. The correlation of the intensity profiles over many pulldowns with the bait intensity profile is reported in the output tables together with the volcano plot-derived significance of the interaction. When assembling the interaction network a threshold is applied to this correlation in order to define an additional class of interactions (Class C), which might not have been found by volcano plot analysis (Classes A and B). This workflow is especially appealing for interaction screens with a large number of bait proteins.

### IMPORTING, CURATING AND PROBING LARGE-SCALE PPI NETWORKS

While protein interaction screens can uncover novel or condition-specific interactions, a wealth of detected and predicted interactions are already stored in protein-protein interaction databases^28^. Analyzing large-scale PPI networks jointly with other omics data has great promise. However, a major obstacle to performing systems-level analysis on these large-scale networks are lacking easy-to-use software solutions to transparently handle the processing and analysis of these networks. Many studies under-utilize the existing resources and mostly report the interactions of a single protein as an after-thought. In the following, we introduce the new network capabilities of Perseus to assemble, filter, and understand large-scale PPI networks, which lay the foundation for any network analysis.

The first task is assembling a high confidence interaction network. Many databases, such as STRING^13^, BioGRID^14^ or HIPPIE^29^, allow researchers to download all interactions in a tabular format, which can be easily loaded into Perseus, even with sizes of up to few millions of interactions. Supplementary information on the interactions such as, but not limited to, the interaction type or a measure of confidence, remain available at each step in the subsequent data analysis. With this information, generalized interaction networks, such as STRING can be filtered by interaction type to generate a physical interaction network. Confidence measures often integrate diverse knowledge into a single score, derived from how often, and by which experimental technique, an interaction was detected, combined with more abstract measures, such as co-expression and literature co-occurrence of the interaction partners^13^. There are two approaches for interaction confidence aware network analysis (Fig. 5a). Applying a cutoff to the confidence score removes low-confidence interactions from the network, which is especially useful when applying methods that treat all interactions equally. The cutoff can be chosen according to the confidence score distribution and the targeted network size (Fig. 5b). Other methods operate on weighted networks and distinguish between interactions with high or low confidence. In this case the confidence scores can be used as an edge weight. In addition to static confidence scores, one can devise dynamic confidence scores from experimental data which reflect e.g. changes in abundance or localization of any of the interactors.

**Figure 5.**
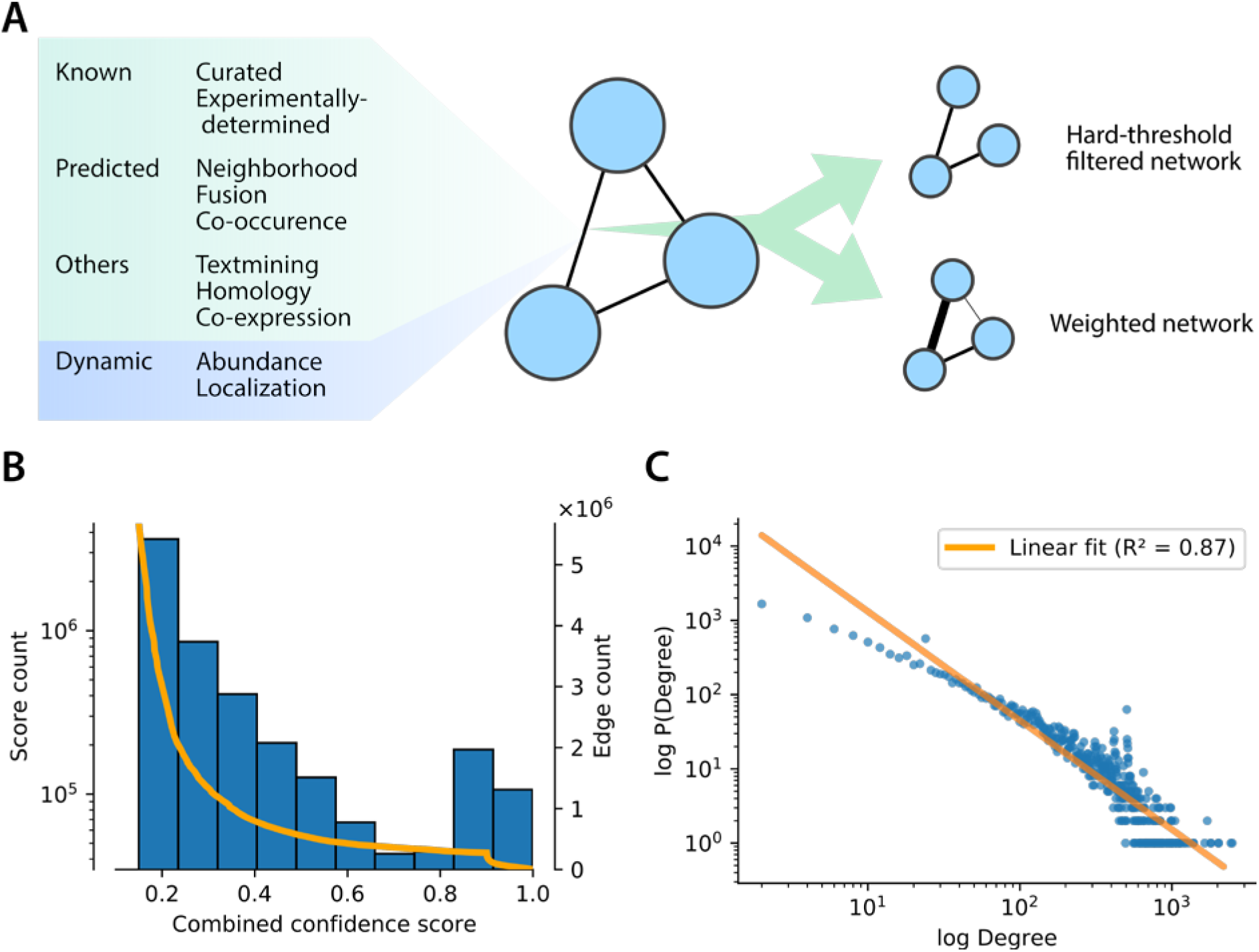
Handling large-scale protein interaction databases in Perseus. a Interactions in PPI databases are often annotated with confidence scores derived from various sources. Static confidence scores can combine experimental evidence and predictions of physical interaction as well as non-physical interactions between the proteins. Confidences can be adjusted dynamically based on condition-specific data to better represent the changed wiring. In any analysis a high-confidence network can be obtained by removing edges below a given hard threshold. Many analysis can directly utilize confidence scores as so-called edge weights, thereby allowing for the inclusion of lower-confidence interactions. b Histogram of the combined confidence score from the human STRING PPI network. Superimposed in orange is the number of interactions in the filtered network if the edges with scores lower than the current value were removed. Filtering out low confidence edges leads to a significant reduction in the number of edges in the final network. c Log-log plot of the node degree against the degree frequency generated from the human STRING PPI network. The R^A^2 value of the linear fit (orange) to these data represents the scale-free fit index.

A deeper understanding of the network requires a different perspective in addition to the interaction-centric view. Any list of interactions can be converted into a network collection with a single click. A dedicated set of network-specific processing activities are now available. While processing the list of interactions the focus remains on the edges of the network. In the network view, the focus is shifted to the nodes. With the powerful identifier and data mapping mechanisms in Perseus, nodes are easily annotated with various annotations, such as gene ontology^30^ (GO), or quantitative proteomics data. Any annotation can be subsequently used to filter the nodes of the network. One could, for example, extract a subnetwork of proteins associated with a specific GO category and their interactions from the large-scale network. Using the data mapping from e.g. deep proteomes of specific cell-lines or tissues, condition-specific subnetworks can be created.

Further understanding is gained by studying the intrinsic properties of networks. By calculating node degrees, corresponding to the number of neighbors of each node in the network, hub nodes can be distinguished from peripheral nodes. By analyzing the distribution of the node degrees in the network, global network properties, such as approximate scale-freeness^31^ of the topology can be identified (Fig. 5c). Furthermore, intrinsic local network properties, like the node degree can be correlated with biological properties derived from protein annotations or experimental data. The proper construction of large-scale interaction networks and understanding their basic properties are central to the successful application of more specialized analyses such as the integration of such networks with PTM data.

### NETWORK ANALYSIS OF PTM DATA

The MS-based study of PTMs is nowadays possible on a global scale for several types of modifications. The best known example is MS-based phosphoproteomics^32^, which is a powerful tool for interrogating signaling events on a large scale. However, drawing conclusions directly from phosphorylation changes is challenging, due to the mostly missing functional information on the inhibitory or excitatory action of a specific protein phosphorylation at a specific site. Network-based approaches for the analysis of phosphorylation data derive functional information on protein-level by interrogating the phosphorylation changes observed in the network neighborhood^33-35^.

We implemented the popular kinase-substrate enrichment analysis^34^ (KSEA) tool for predicting kinase activities in Perseus. Site-specific kinase-substrate networks (Fig. 6a) assign kinases to the experimentally observed phosphorylation sites. The core of the analysis is the calculation of a series of scores (mean, enrichment, Z-score, p-value, q-value) for each kinase, based on the quantitative phosphorylation changes of its substrates. These predicted kinase activities can be analyzed further to find e.g. differentially activated kinases or pathways. KSEA most often utilizes the curated kinase-substrate network from the PhosphoSitePlus database^17,36,37^. In order to extend the coverage of the network and thereby allow for the utilization of a larger fraction of the experimental data, the network can be supplemented with predicted kinase-substrate interactions from tools such as NetworKIN^38,39^, or with low-specificity interactions derived from kinase target sequence motifs.

PHOTON^33^, now implemented in Perseus, is an alternative approach to KSEA that calculates more broadly defined signaling functionality scores for any protein, rather than activities for kinases only. A data-annotated large-scale PPI network now serves as the input (Fig. 6a). The resulting signaling functionality scores for each experimental condition are based on the observed phosphorylation in the neighborhood of each proteins and are assigned a significance by a permutation-based FDR scheme. The scores can either be analyzed directly, to find proteins and pathways with e.g. differentially changing signaling functionality or utilized in a second step of the PHOTON pipeline, in which signaling pathways are automatically reconstructed from the large-scale network, that connect the proteins with significant signaling functionality.

**Figure 6.**
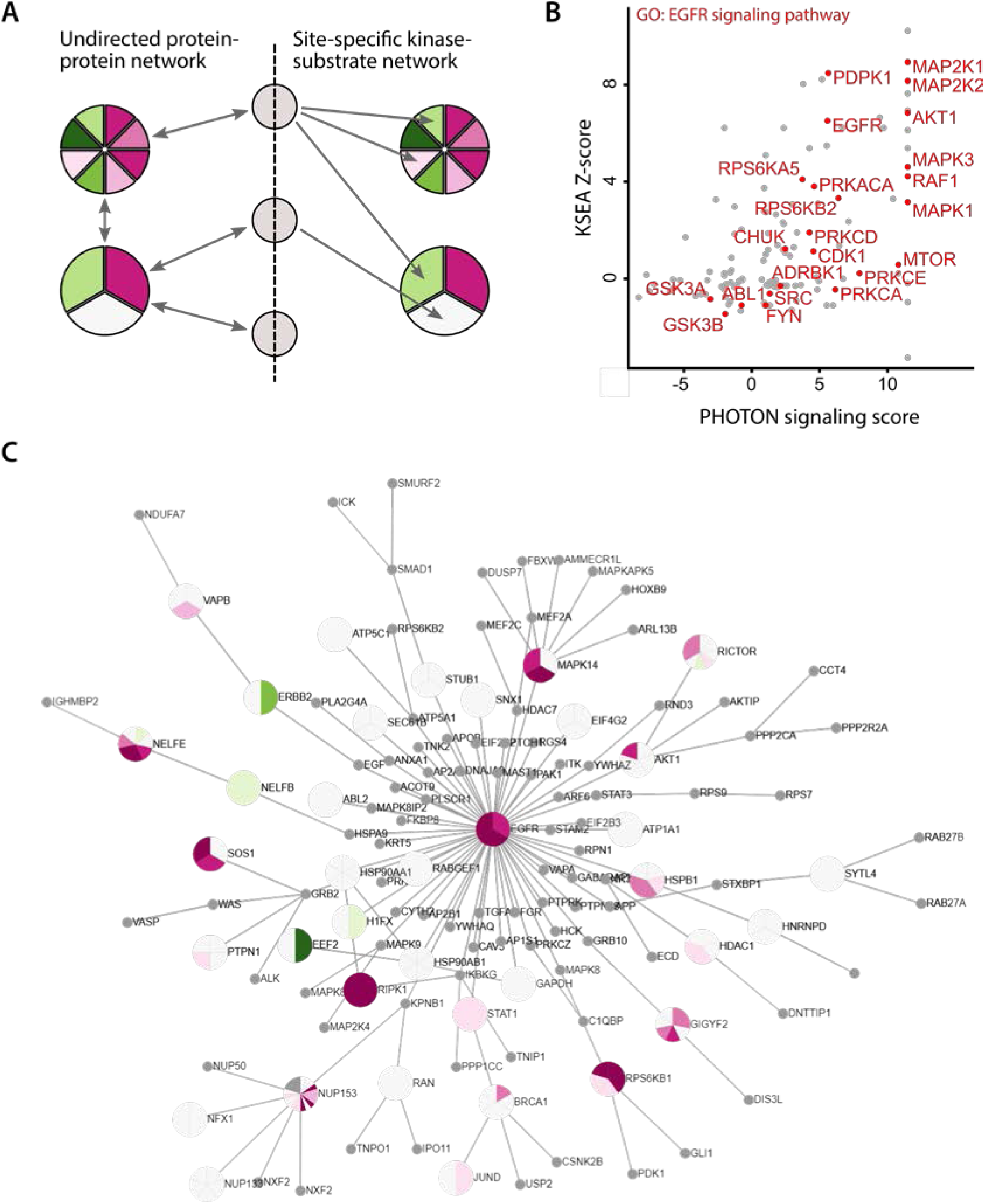
Network analysis of an EGF stimulation phospho-proteomics study. a Comparison of network topologies used for the analysis of phospho-proteomic data. Nodes in the network are represented as grey circles or pie-charts where each slice represents the observed phosphorylation changes at a specific site on the protein. Physical protein-protein interactions (left side) are present between all classes of proteins and are by definition undirected. In order to capture the enzymatic action of kinases more accurately, directed interactions (right side) from kinase to substrate are defined in a site-specific manner. b KSEA Z-score and PHOTON signaling functionality scores derived from phospho-proteomic data measured after EGF stimulation (Supplementary Table 2) only weakly correlate to each other (Pearson correlation 0.52). Kinases annotated in GO with the term ‘Epidermal growth factor receptor signaling pathway’ are highlighted in red. Both methods assign high scores to central members of the expected pathway. c Signaling network reconstructed by PHOTON from the 100 highest scoring proteins anchored at EGF. The interactive visualization has an automatic layout and phosphorylation data overlay.

The Perseus network module allows for performing both KSEA, and PHOTON analysis on the same experimental data^33^ and a choice of networks^17,29^. Due to the differences in the utilized methodologies and the chosen networks, resulting scores will differ, but are easily compared with the analyses and visualizations provided by Perseus (Fig. 6b). Both tools support the analysis of datasets with multiple conditions, effectively transforming the peptide-level phosphorylation data into protein-level scores. The entire well established toolset for the analysis of protein quantification data can be applied to these scores, including hierarchical clustering, enrichment analysis^33^ and time-series analysis^40^.

We implemented an interactive visualization of kinase-substrate networks directly in Perseus (Fig. 6c) using the cytoscape.js library^25^. The visualization allows for the joint visual inspection of the networks, e.g. subnetworks reconstructed by PHOTON, and the quantitative phosphorylation data. Browsing the quantitative phosphorylation in a reduced and highly structured network view while also considering the signaling functionality scores, allows for the generation of hypotheses that explain the signal transduction mechanistically.

### CO-EXPRESSION CLUSTERING AND CLINICAL DATA

When performing co-expression analysis, the correlation matrix between the proteins in the dataset describes a fully connected, weighted network, in which the weight on each edge denotes the correlation between the two proteins (Fig. 7a). Hence, the actual network usually remains implicit. A hierarchical clustering of the co-expression network can utilize the network neighborhood of each protein and integrate it into the similarity calculation^41^. The cluster dendrogram and the detected co-expression modules are then transferred back to the original data where their interpretation is equivalent to ordinary hierarchical clustering. In addition to the clustering, a representative expression profile for each of the clusters is generated, which is termed eigengene. This highly reduced view on the data can be correlated with clinical or phenotype data and clustered to gain a better understanding of the behavior of the detected cluster (Fig. 7b). The described co-expression analysis is available in Perseus through the R language interface provided by PluginInterop which interfaces directly with the established WGCNA library^42^.

**Figure 7.**
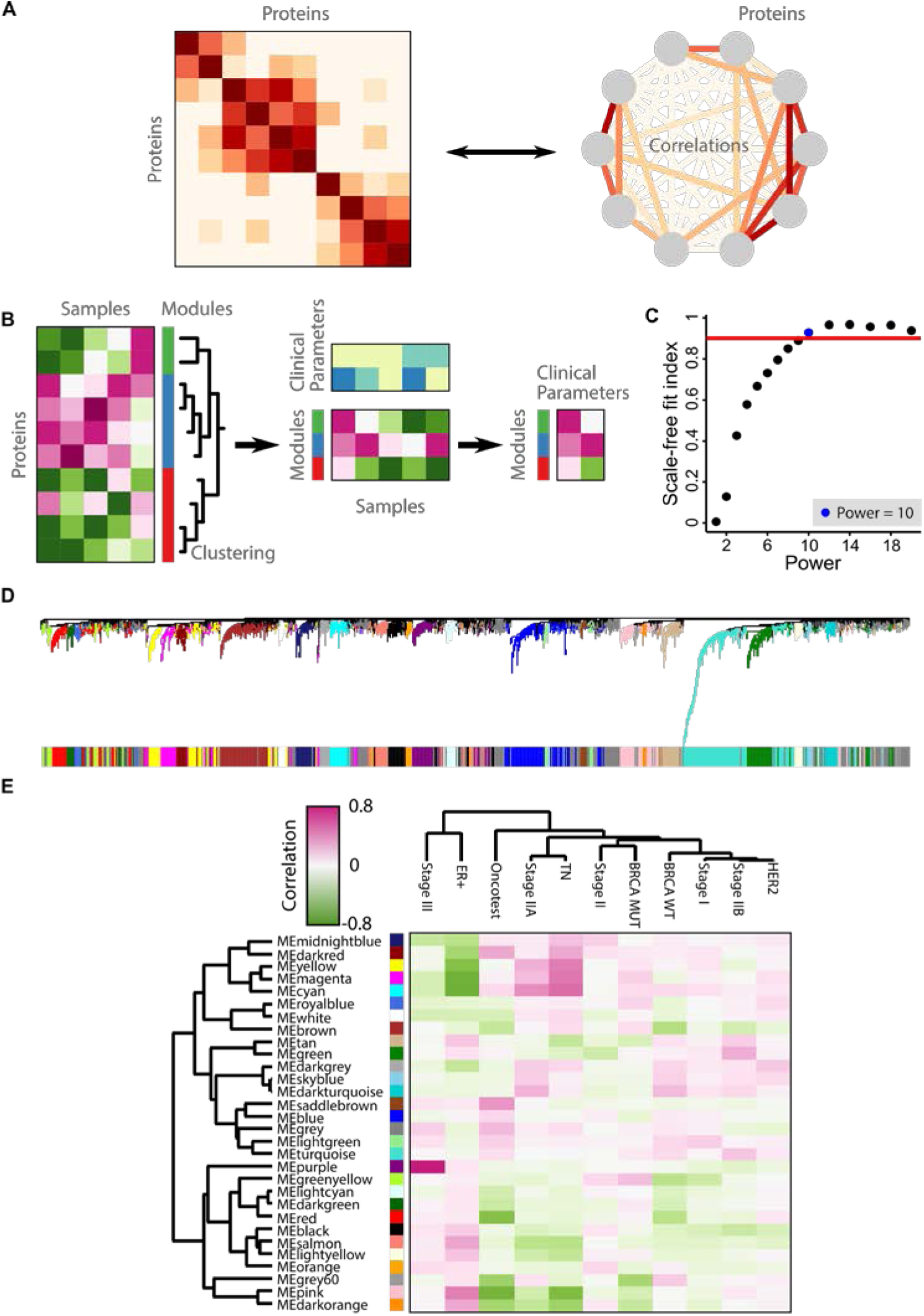
Co-expression network analysis on clinical data. a Any correlation matrix can be interpreted as a fully-connected network with edge weights corresponding to the correlation between the proteins. Hierarchical clustering of the correlation matrix can utilize a network-based distance function. b Co-expression clustering and identified co-expression modules annotate the original expression matrix. Phenotype data can be correlated with representative co-expression module profiles and provide a high-level interpretation of the modules. c Parameter selection of the power parameter for the Yanovich et al.^43^ dataset (Supplementary Table 3 and Online Methods). The lowest power reaching close to a high scale-free fit index of 0.9 (red line) was selected. d Co-expression cluster dendrogram and assigned modules. e Correlation heat map between module eigengenes and clinical parameters.

We applied the WGCNA co-expression analysis to parts of a cancer proteomics dataset^43^, following the recommended workflow^7^ from within Perseus. Bi-weight midcorrelation, a robust alternative to Pearson correlation, was chosen to calculate correlations between all pairs of proteins. In order to obtain a scale-free co-expression network, a power parameter of 10 was selected (Fig. 7c), leading to an approximately scale-free network with a scale-free fit index of 0.9. Hierarchical clustering of the co-expression network identified 30 modules (Fig. 7d). The representative expression profiles of each of the modules, as provided by the corresponding module eigengene, were correlated with the available clinical annotations. This high-level overview over the data was then visualized in a heatmap (Fig. 7e). Several modules showed high correlations with specific clinical annotations. The magenta module showed high correlation with the triple-negative subtype (TN) and was highly enriched for the ‘interferon-gamma-mediated signaling pathway’ GO category. The top module hub genes with kME > 0.8 were GBP1, TAP1, TAPBP, HLA-A, TAP2, STAT1, and EML4. The purple module showed high correlation with Stage III, but and in depth look at the co-expression clustering heatmap revealed the module to be dominated by a single patient, limiting the validity of the module.

### SOFTWARE IMPLEMENTATION, DOWNLOAD AND MAINTENANCE

The Perseus network module PerseusNet is implemented in the C# programming language using Visual Studio 2017, like the whole Perseus software. PerseusNet is distributed with Perseus by default and can be downloaded from http://www.perseus-framework.org. The current version, which is described in this manuscript, is 1.6.2.3. The PluginInterop and PHOTON plugins are also included in the standard download. In the current release, it is recommended to use Windows as operating system, although Linux support is underway, realized in the same way as for the MaxQuant software^44,45^, by ensuring Mono compatibility. A plugin API enables external programmers to extend the functionality of PerseusNet and Perseus in general, by programming their own workflow activities. Plugin extensions by the user community will be linked from the plugin store at http://www.coxdocs.org/doku.php?id=perseus:user:plugins:store upon request. Context-specific documentation is linked from each activity (Supplementary Fig. 3). Step-by-step guides for the integration of external tools, such as Python or R, that have to be installed and configured separately from the main Perseus software, are available online^8^. A help forum for Perseus and PerseusNet is available at https://groups.google.com/group/perseus-list. Bugs that are reproducible in the latest available software version should be reported at https://maxquant.myjetbrains.com/youtrack.

## DISCUSSION

We introduced PerseusNet, the network analysis extension for the Perseus software. It enables proteomics researchers to perform most network analysis by themselves. PerseusNet is highly extensible through a plugin API and its extension to R and Python, which allows for the incorporation of a plethora of existing scripts and programs from the network community. We envision that large part of the future programming will not be done by local developers, but by the global community through the plugin API. Programmers can release their plugins under licenses of their choice.

We have implemented powerful proteomics-specific activities for AP-MS network generation and PTM-related network analysis, presumably the two main applications for networking in proteomics. We plan to extend PerseusNet in the near future by activities from other proteomics sub-domains, as interaction determination by protein correlation profiling^46^ and large scale network generation from cross-linking experiments on whole cell lysates^47^.

## ONLINE METHODS

### CREATING INTERACTION NETWORKS FROM PULL-DOWN EXPERIMENTS

We created an interaction network from a pull-down screen^27^ (Fig. 4). First, .RAW files were obtained from PRIDE (PXD003758) and processed with MaxQuant version 1.6.2.10. Mouse protein sequences were downloaded from UniProt (release 2017_07). Parameters ‘matching between runs’ and ‘LFQ’ were selected in addition to the default parameters. Downstream analysis of the ‘proteinGroups.txt’ output table was performed in Perseus. Columns for baits Eed, Ring1b, and Bap1 and their controls in the ESC and NPC cell lines were selected and log transformed. Quantitative profiles were filtered for missing values, and were filtered independently for each of the bait control pairs, retaining only proteins that were quantified in all three replicates of either the bait, or control, pull down. Missing values were imputed (width 0.3, down shift 1.8) before combining the tables, resulting in a total list of 2995 proteins on which the multi-volcano analysis was performed (Supplementary Table 1). The s0 and FDR parameters for Class A (s0=1, FDR=0.01%) and Class B (s0=1, FDR=0.2%) were chosen by visual inspection, aiming for a low number of significantly depleted proteins in any of the experiments. The interaction network was created by connecting significantly enriched prey proteins to their baits. Edges representing known protein complex interactions were annotated in the network. Due to missing mouse CORUM annotations for any of the baits, mouse CORUM annotations were obtained by mapping between mouse and human homologues as listed in the MGI^48^ database. The resulting annotated network was then visualized in Perseus.

#### Approximately scale-free topology of the STRING interaction network

In order to investigate the topology of a large scale interaction network (Fig. 5), we first downloaded the human STRING interaction network (v10.5) from the website. After filtering for high confidence interactions (>0.9), the scale-free fit index was calculated according to reference^42^. Node degrees were calculated and plotted against their frequency distribution on a log-log scale. The R^^^2 of a linear fit to the log-log space represents the scale-free fit index.

### NETWORK ANALYSIS OF A PHOSPHO-PROTEOMIC DATASET OF EGF STIMULATION

Two separate analyses, PHOTON and KSEA, were performed on the same experimental dataset^33^ (Supplementary Table 2) and compared (Fig. 6). Log2 fold-changes for EGF from two replicates were averaged. For PHOTON analysis, we first generated a high-confidence PPI network. We downloaded all interactions from HIPPIE and filtered them for high confidence interactions (confidence > 0.72) and additionally removing high-degree nodes (degree < 700). The experimental data was mapped from UniProt to Entrez GeneIDs and subsequently used to annotate the nodes of the network. We then performed PHOTON analysis with adjusted default parameters. Network reconstruction with ANAT was enabled with the 100 highest scoring proteins and EGF anchor (GeneID 1950). Additionally, we increased the number of permutations to 100,000. The KSEA analysis was performed on the human site-specific kinase-substrate network from PhosphositePlus^17^. Data and network were matched based on UniProt identifiers. The resulting KSEA Z-scores and PHOTON signaling functionality scores were plotted against each other in Perseus. Proteins annotated with the GO category ‘Epidermal growth factor receptor signaling pathway’ (GO:0007173), were highlighted in red.

#### Co-expression analysis of a clinical proteomics dataset

We applied the co-expression network analysis workflow to a clinical proteomics dataset (Fig. 7). Protein quantification data and clinical annotation were obtained from Yanovic et al.^43^. SILAC ratios were first transformed to log(light/heavy). The dataset was filtered for the 43 patients unique to Yanovich et al. Using global hierarchical clustering of the patients, 4 outlier samples were identified and removed from the dataset. Additionally, proteins with less than 70% valid values were removed from the dataset and the resulting patient profiles were Z-scored (Supplementary Table 3). The power parameter for the co-expression analysis was selected using the ‘oft-threshold’ activity. Using a signed network and the biweight midcorrelation, the power 10 was the lowest to a have scale-free fit index of more than 0.9 (Figure 7c). Proteins where then subjected to co-expression clustering (Figure 7d) and 30 co-expression modules were identified. The eigengene of each co-expression module was correlated with the provided clinical data using Pearson correlation and clustered using hierarchical clustering (Figure 7e).

### PLUGININTEROP PROVIDES A CENTRAL ENTRY-POINT FOR ALL EXTERNAL PLUGINS

The PluginInterop project is written in the C# programming language and implements several Perseus plugin APIs. For users it provides a number of activities in Perseus for executing script files written in the Python and R languages. Upon selection of any of these activities, users will be prompted with a parameter window, allowing them to pass additional arguments to the script and requiring them to specify the executable that should be used for processing. Since Perseus does not include an installation of Python or R, users will have to install those and any other dependencies separately. PluginInterop aids the user by trying to automatically detect an existing installation and provide meaningful error messages in case of missing dependencies. Developers can additionally leverage the functionality implemented in PluginInterop as a basis for parametrized scripts. In general, developers are free to choose which external scripting language or program they would like to utilize. We found the R and Python scripting languages to be most useful, which is why we provide two companion libraries ‘perseuspy’ and ‘ PerseusR’ to be used alongside PluginInterop. These libraries aid the communication between Perseus and the script.

The communication between Perseus and external scripts is straightforward and is easily implemented for any tools of choice. In short, Perseus will persist all necessary data to the hard-drive and call the specified tool with specific command-line arguments. The first arguments contain all the parameters specified by the user, per choice of the developer, either in an XML format, or simply separated by spaces. Secondly, the input data from the workflow is saved to a temporary location which is passed to the script. The final arguments specify the expected location of the output data. The external process can provide status and progress updates to the user, as well as detailed error reporting by printing to stdout/stderr and indicating success or failure through the exit code. Once the process exits, Perseus will parse the output data for its expected location and insert it to the workflow. Any step in the pipeline is customizable for advanced scenarios, such as custom data formats.

The PluginInterop binary is automatically included in the latest Perseus version. The source code was published under the permissive, open-source MIT license, on Github^9^. The website also provides more information on how to develop plugins, including a video demonstration. The plugins presented in this manuscript are all developed on top of PluginInterop and the perseuspy and PerseusR companion libraries.

### LIBRARY SUPPORT FOR SCRIPTING LANGUAGES

We implemented libraries in R and Python, which facilitate the interoperability of Perseus with external scripting languages. The main aim of these libraries is to map the data structures of Perseus to a counterpart native to the external language. Developers proficient in these languages will be more comfortable and productive with these native data structures. The largest benefit comes from the resulting integration with the existing data science ecosystem, all now available to Perseus plugin developers.

The ‘perseuspy’ module provides data mappings for the Python language. The Perseus expression matrix is mapped the ‘DataFrame’ object of the popular ‘pandas’ module, which is tightly integrated with ‘numpy’, the de-facto standard for numerical computations in Python. The Perseus network collection data-type maps to a list of networks from the ‘networkx’ package. It features a variety of graph algorithms and interfaces well with other modules, due to its usage of standard Python dictionaries. ‘perseuspy’ is distributed via The Python Package Index (PyPI), allowing for easy installation of the module for developers and users alike. The code of ‘persuspy’ is published under the permissive, open-source MIT license, and available alongside usage examples and more information on https://github.com/jdrudolph/perseuspy.

For the R language, we implemented the ‘PerseusR’ package. It provides a mapping of the Perseus expression matrix to a custom wrapper class around the R ‘data.frame’ object. The wrapping was necessary to represent Perseus-specific information such as annotation rows. Alternatively, developers can load data as a Bioconductor ‘expressionSet’ object which enables the interface with the entire Bioconductors bioinformatics suite. Currently there is no support for network collections in ‘PerseusR’, but we plan to implement it in the near future. ‘PerseusR’ is also published under the MIT license and its code is available on https://github.com/jdrudolph/PerseusR. Currently ‘PerseusR’ is easily installed directly from the website. Due to the lengthy submission process, ‘PerseusR’ will be uploaded to CRAN at a later point in time.

### IMPLEMENTATION OF PLUGINPHOTON

We implemented a Perseus plugin for the PHOTON tool on top of the functionality provided by PluginInterop and perseuspy. PHOTON was previously capable to run only a single experiment at a time with a fixed human protein-protein interaction network. We expanded its implementation to allow for parallel processing of any number of experiments on any network. These changes make large datasets from any species directly amenable to PHOTON analysis. PluginPHOTON is published under the MIT licence, its code is available on https://github.com/jdrudolph/photon, and it is included in the latest Perseus release.

### IMPLEMENTATION OF PLUGINCOEXPRESSION

We implemented parts of the WGCNA pipeline as a Perseus plugin. PluginCoExpression provides access to the WGCNA functions implemented in the R language via PluginInterop and PerseusR.

#### Implementation of KSEA in Perseus

KSEA analysis was implemented in Perseus and tested for correctness against the reference implementation.

## ACKNOWLEDGEMENTS

We thank J. Sebastian Paez and Sung-Huan Yu for contributing to PerseusR, and Caroline Friedel and Tamar Geiger for helpful discussions. This project has received funding from the European Union′s Horizon 2020 research and innovation programme under grant agreement no. 686547 and from the FP7 grant agreement GA ERC-2012-SyG_318987–ToPAG.

## AUTHOR CONTRIBUTIONS

J.R. and J.C. planned and performed the research, developed the software and wrote the manuscript.

## COMPETING FINANCIAL INTERESTS

The authors declare no competing financial interests.

## Footnotes

1 http://graphml.graphdrawing.org/

2 http://manual.cytoscape.org/en/stable/Supported_Network_File_Formats.html

3 http://jsongraphformat.info/

4 PerseusR, https://github.com/jdrudolph/PerseusR

5 perseuspy, https://github.com/jdrudolph/perseuspy

6 https://github.com/jdrudolph/PluginInterop

7 http://www.peterlangfelder.com/wgcna-resources-on-the-web/

8 https://github.com/jdrudolph/PluginInterop

9 https://github.com/jdrudolph/PluginInterop

